# Comparison of *Escherichia coli* surface attachment methods for single-cell, *in vivo* microscopy

**DOI:** 10.1101/648840

**Authors:** Yao-Kuan Wang, Ekaterina Krasnopeeva, Ssu-Yuan Lin, Fan Bai, Teuta Pilizota, Chien-Jung Lo

## Abstract

For *in vivo*, single-cell imaging bacterial cells are commonly immobilised via physical confinement or surface attachment. Different surface attachment methods have been used both for atomic force and optical microscopy (including super resolution), and some have been reported to affect bacterial physiology. However, a systematic comparison of the effects these attachment methods have on the bacterial physiology is lacking. Here we present such a comparison for bacterium *Escherichia coli*, and assess the growth rate, size and intracellular pH of cells growing attached to different, commonly used, surfaces. We demonstrate that *E. coli* grow at the same rate, length and internal pH on all the tested surfaces when in the same growth medium. The result suggests that tested attachment methods can be used interchangeably when studying *E. coli* physiology.

## Introduction

Microscopy has been a powerful tool for studying biological processes on the cellular level, ever since the first discovery of microorganisms by Antonie van Leeuwenhoek back in 17th century^1^. Recently employed, *in vivo* single-cell imaging allowed scientists to study population diversity^2^, physiology^3^, sub-cellular features^4^, and protein dynamics^5^ in real-time. Single cell imaging of bacteria is particularly dependent on immobilisation, as majority of bacteria are small in size and capable of swimming. Immobilisation methods vary depending on the application, but typically fall into one of the two categories: use of physical confinement or attachment to the surface via specific molecules. The former group includes microfluidic platforms capable of mechanical trapping^6, 7^, where some popular examples include the “mother machine”^8^, CellASIC^9^ or MACS^10^ devices, and porous membranes such as agarose gel pads^2, 11–13^. Physical confinement methods, while higher in throughput, can have drawbacks. For example, agarose gel pads do not allow fast medium exchange, and when mechanically confining bacteria the choice of enclosure dimensions should be done carefully in order to avoid influencing the growth and morphology with mechanical forces^14^. Furthermore, mechanically confined bacteria cannot be used for studies of bacterial motility or energetics via detection of bacteria flagellar motor rotation^15–17^. Chemical attachment methods rely on the interaction of various adhesive molecules, deposited on the cover glass surface, with the cell itself. Adhesion can be a result of electrostatic (polyethylenimine (PEI)^18, 19^, poly-L-lysine (PLL)^15, 17, 19^) or covalent interactions (3-aminopropyltriethoxysilane (APTES)^19^), or a combination, such as with polyphenolic proteins (Cell-Tak^*TM*^)^19^.

Time scales on which researchers perform single-cell experiments vary. For example, scanning methods, like atomic force microscopy (AFM) or confocal laser scanning microscopy (CLSM)^19, 20^, require enough time to probe each point of the sample, and stochastic approaches of super resolution microscopy (e.g. PALM and STORM) use low activation rate of fluorophores to achieve a single fluorophore localisation^20, 21^. Thus, required acquisition time scales vary from milliseconds to minutes^21, 22^, while experiments aimed at the observation of cell growth or slow cellular responses can run from minutes to hours^8, 11, 16, 23, 24^.

Regardless of the time scale, physiology of the studied bacteria should not be affected by the adhesives used for surface attachment. For example, single particle tracking is often performed on surface immobilised cells^5, 25, 26^, and cellular physiology can influence particle diffusion in the cytoplasm, e.g. metabolic “stirring” of the cytoplasm enhances diffusion in size dependant manner^27, 28^, and intracellular pH of yeast has been shown to affect cytoplasm fluidity^29^. Furthermore, concerns have been raised that charged molecules, like PLL that is commonly used for a surface attachment^17, 24, 25, 30, 31^, can perturb membrane potential causing partial or complete membrane depolarisation^32, 33^. Additionally, PLL in high concentrations exhibits antimicrobial properties^34^. Despite of these concerns, PLL has been widely utilized in super-resolution and single molecules tracking applications as it is cheap and easy to use^35, 36^. It is, therefore, important to characterise physiological parameters on different surfaces, and on the time scales relevant for live cell imaging.

In this report we compare a range of immobilisation techniques, including PLL, PEI, Cell-Tak^*TM*^and agarose gel pad, using *Escherichia coli* as a model organism. We measure several physiological traits during growth on the specific surface, including growth rate, size and intracellular pH, and find that tested immobilisation methods do not differ; growth rate and cell size are surface-attachment independent.

## Results

### Immobilisation assays

We test four substrates commonly used for bacteria immobilisation: poly-L-lysine (PLL)^15, 33^, polyethylenimine (PEI)^18, 37^, Cell-Tak*™* ^19^, 3-aminopropyltriethoxysilane (APTES)^19^ and agarose gel^22^.

PLL and PEI are cationic polymers, which can electrostatically interact with negative charges on the outer surface of the cell^38^. Several PLL coating protocols have been reported, which we here refer to as “in-chamber”^15^, “rinsed”^33^, and “air-dried”^33^ methods. “In-chamber” PLL coating is the most standard for bacterial flagellar motor experiments commonly known as bead assay^15, 39–41^. In this protocol PLL solution is flushed into an uncoated glass flow-chamber for no longer than 15 s followed by thorough washing with the excessive volume of growth medium (~ 25 times the flow-chamber volume). In the “rinsed” method lower PLL concentration and longer incubation time (min) are used to cover the entire surface of the coverslip by immersing it in the PLL solution^33^ and subsequent washing. “Air-dried” method is similar to the “rinsed”, with an addition of complete drying the PLL solution on the coated surface for over an hour before washing. For our detailed coating protocols see *Materials and Methods*.

Cell-Tak*™* is a commercially available adhesive extracted from marine mussel, *Mytilus edulis*. It’s a component of byssus, a bundle of filaments mussels secrete to anchor themselves to solid surfaces^42, 43^. Characterising and mimicking the adhesive chemistry of mussel byssus is an active area of research^44^. What we know thus far, is that it involves bidentate and covalent interactions, protein coacervation, intrinsic protein-protein binding as well as metal chelation^44^. We use the manufacturer coating protocol as described in *Materials and Methods*.

APTES is a common choice for salinisation of microfluidic channels^45^, and has also been employed as an attachment agent for the AFM imaging^19^. We coat coverslips by incubating them in a 2% solution of APTES for 2 h followed by extensive washing with water and acetone as described in *Materials and Methods*.

As a control we grow bacteria on the agarose gel pad with no chemical adhesives (see *Materials and Methods* for details).

### Growth rate and morphology of bacteria do not dependent on the surface attachment method

To estimate the effect of different immobilisation assays on bacterial physiology we examine the growth rate, size and division accuracy of individual cells growing on each specific surface. The growth regulation of bacteria has been attracting researchers attention since 1950s. As a result, steady state population growth rates at different carbon sources and temperatures^47, 48^, the nature of single cell growth and its connection with the population growth^46, 49–51^, the relationship between cell size and growth rate^47, 52^, as well as the accuracy and robustness of cell division, have all been established^53–55^. The population growth rate (*λ*) was found to depend on the nutritional composition of the media, and was slower at lower temperatures^47, 56^. Here *λ* refers to the cell population growing exponentially, also called balanced growth or steady state growth, so that *N*(*t*) = *N*_0_⋅*e*^*λ t*^, where *N*(*t*) is the number of cells at a given time point *t*. As the number of cells in the population doubles with a specific doubling time *t*_*D*_, *N*(*t*) can be expressed as 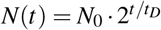, and *λ* = *ln*(2)/*t*_*D*_. Individual cells within the population were also found to grow exponentially, specifically in mass and length^46, 49^. Thus, *L*(*t*) = *L*_*b*_ ⋅ *e*^*bt*^, where *L*_*b*_ is the cell length at birth and *b* the growth rate of an individual cell, which is equal to *b* = *ln*(*L*_*d*_/*L*_*b*_)*t*_*D*_ ^50, 51^. *L*_*d*_ is a cell length at division and in *E. coli* it is equal to two lengths of the cell at birth^53–55^. Therefore, to a good approximation *λ* = *b* (if noise is taken into the account the population growth rate is slightly lower when compared to *b*^50, 51^). In summary, from previous work we expect that if the specific surface attachment method does not alter cells’ physiology in comparison to planktonic growth, *b* will be equal to *λ* (for a given medium and temperature), the cell size should be a function of growth rate and the cells should divide in half.

To obtain *b* and cell size we monitor bacteria between the first and second divisions (*G*_1_ and *G*_2_, see also *Materials and Methods* and Figure 1, top) using optical microscopy. Phase-contrast images of the bacteria are taken every 5 min and cells’ length and width (*W*) are extracted as described in *Materials and Methods* and Figure 1A. Prior to imaging cells are grown in a flask at 37°C and later kept at 23°C during flow chamber preparation for ~ 40 min (see also *Materials and Methods*). Bacterial growth assay is then performed at 23°C in a flow chamber. SI Figure 1 shows growth rates and bacterial length versus time from the beginning of the imaging (*G*_0_ to *G*_2_ as defined in Figure 1, top), indicating that by the time *G*_1_ is reached, cells’ growth rate is approximately constant. Thus, unless explicitly stated, from here on we combine *b*, *L* and *W* from *G*_1_ and *G*_2_. During imaging fresh oxygenated medium is continuously supplied to the cells.

**Figure 1.**
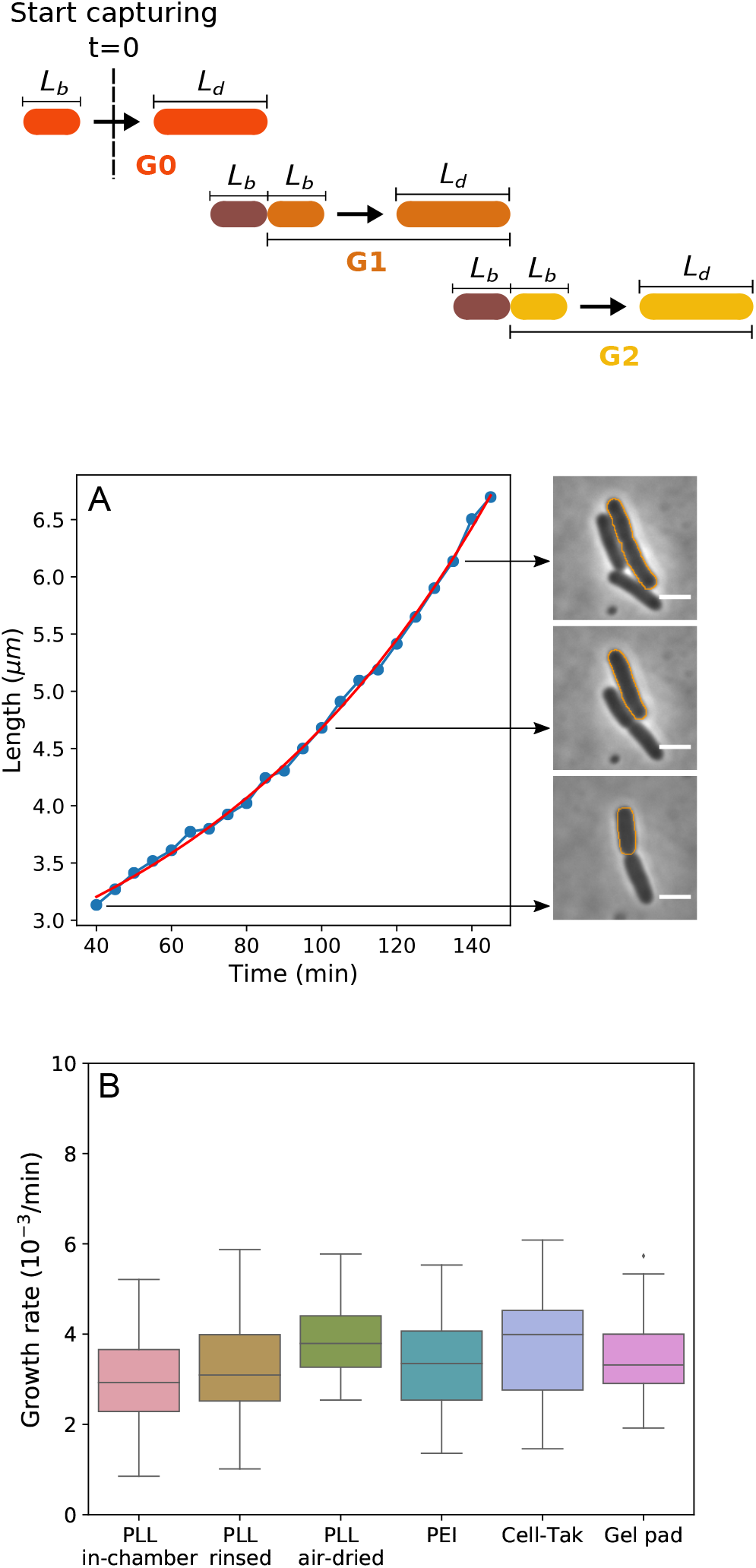
Bacterial growth rate stays constant when cells are attached to the surface with different attachment methods.Top, a cartoon depicting our imaging protocol. *G*_0_ are the cells observed upon commencing the imaging to the first visible division. Subsequent two generations (*G*_1_ and *G*_2_) are used to obtain our results (unless otherwise stated). (**A**) The single cell length is measured using phase contrast imaging over a complete first generation (G1) cell cycle. Red line shows the fit to a single exponential function from which we obtain the single cell growth rate^46^. The diagram above shows the definition of different generations. We call the generation zero (G0) cells that were deposited onto the surface after their growth on the flask. We start our observation in the middle of the G0 growth cycle. G0 is followed by the generation one (G1) that lasts from the first observed division to the second, and so on. Panel on the right shows the phase contrast images of the tracked cell (marked with the orange outline) at the indicated time points. Scale bar is 2 µm. (**B**) Comparison of average growth rates on different surfaces shows no significant difference between all tested immobilisation methods in MM9 medium. The growth rates were measured at 23°C. Number of cells analysed for each attachment method is: N=44, 104, 28, 81, 48, and 43 for “PLL in-chamber”, “PLL rinsed”, “PLL air-dried”, PEI, Cell-Tak and gel pad respectively.

Figure 1B shows that single cell growth rates of the cells grown on different surfaces are independent of the surface attachment and in agreement with population growth rates measured by us (SI Figure 5 and Tables 1 and SI 2), and reported previously^47, 48^ (for comparison between *G*_1_ and *G*_2_ see SI Figure 2). As expected, the growth rate is medium dependant and increases when we move from the MM9 medium (see *Materials and Methods*) to the richer LB medium (SI Figure 4A).

**Table 1.** Summary of the experimentally measured variables. Experiments were performed in LB only in the gel pad. Results obtained for *G*_1_ and *G*_2_ phase on each different surfaces (including gel pad) are averaged.

| Medium | $b(\text{min}^{-1})$ | $\lambda(\text{min}^{-1})$ | $L_b(\mu\text{m})$ | $L_d(\mu\text{m})$ | $W(\mu\text{m})$ | $L_b/L_d$ | $L_{\text{daughter}}/L_{\text{mother}}$ | $pH_{\text{initial}}$ | $pH_{\text{final}}$ |
| --- | --- | --- | --- | --- | --- | --- | --- | --- | --- |
| MM9 | $0.0033 \pm 0.0010$ | 0.0032 | $2.61 \pm 0.31$ | $4.35 \pm 0.49$ | $0.94 \pm 0.04$ | $0.60 \pm 0.07$ | $0.50 \pm 0.03$ | $8.39 \pm 0.33$ | $7.82 \pm 0.24$ |
| LB | $0.0083 \pm 0.0011$ | 0.0080 | $3.31 \pm 0.54$ | $5.72 \pm 0.95$ | $1.03 \pm 0.07$ | $0.58 \pm 0.07$ | $0.50 \pm 0.02$ | $8.29 \pm 0.16$ | $8.43 \pm 0.16$ |

In Figure 2A, 2B and Table 1 we analyse the length of individual cells at the beginning (length at birth, *L*_*d*_) and at the end of the growth cycle (length at division, *L*_*d*_), as well as cell width (*W*). Similarly to the growth rates, *L*_*b*_, *L*_*d*_, and *W* are not dependant on the immobilisation method, but do change with the growth media, SI Figure 4B and 4C. However, cells growing on the surface, while growing at the same rate as planktonic cells, divide earlier and become shorter, and equally so on all the surfaces, Figure 2C. Thus, planktonic *t*_*D*_ = *ln*(2)*/λ* and *t*_*D*_ = *ln*(*L*_*d*_/*L*_*b*_)*/b* of cells growing on the surface are not the same despite the same growth rates (Figure SI Figure 3 shows *t*_*D*_ = *ln*(*L*_*d*_/*L*_*b*_)*/b*). When growing on different surfaces cells still divide in half, as shown in Figure 2D and SI Figure 2C, where we plot the length of the mother and daughter cells (*L*_*daughter*_/*L*_*mother*_). We note that this ratio is between two generations, whereas *L*_*b*_/*L*_*d*_ is within one generation (as defined in Figure 1). We summarise our and previous population growth rates in SI Table 2 and all other experimentally measured variables in Table 1.

**Figure 2.**
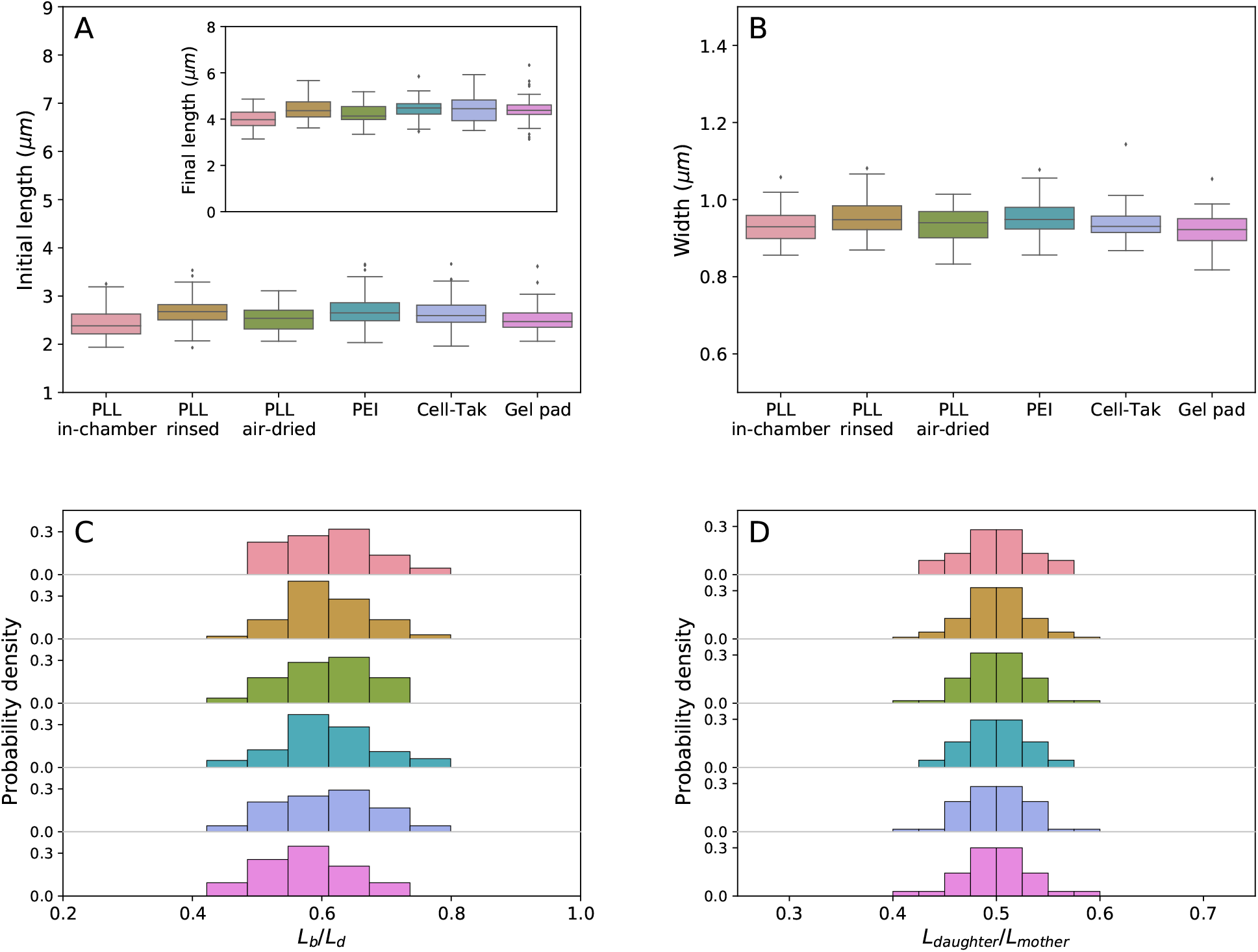
Morphology of bacteria is not influenced by the attachment method, but does change when cells grow on a surface. (**A**) Cell length at birth (*L*_*b*_) of cells immobilised on different surfaces. The inset shows the length at division. (**B**) Width distribution of cells attached to the surface with various coatings remains constant. (**C**) Probability density of length ratio (*L*_*b*_/*L*_*d*_) and (**D**) (*L*_*daughter*_/*L*_*mother*_) remains unchanged on all tested surfaces. Colour coding is as given in A and B.

Cells can grow and divide while attached to the APTES coated surface. However, we were unable to calculate the growth rate, shape or the internal pH due to poor quality of attachment (most of the cells on APTES surface detached withing the first generation). Time lapse video demonstrating cell growth and attachment on APTES coated surface is shown in SI Video 1, in comparison to growth on other surfaces shown in SI Video 2 to 7.

### Intracellular pH during growth on the surface does not depend on the method of attachment

Neutrophilic bacteria maintain their cytoplasmic pH withing a narrow range (termed pH homeostasis). For example, *E. coli* can survive in a range of external pHs, starting as low as pH ~ 2 in the human stomach and up to pH ~ 9 at the pancreatic duct, while maintaining internal pH in a relatively narrow range of 7-8^57–62^. Cytoplasmic pH plays an important role in cellular energetics as the difference between cytoplasmic and extracellular pH contributes to the electrochemical gradient of protons (so called proton motive force^63^), as well as influences protein stability and an enzymatic activity in the cell^64^. However, cytoplasmic pH can change when cells are subjected to an external stress, such as acid or osmotic shocks^61, 65, 66^. Furthermore, for some species acidification of the cytoplasm has been shown to be related to pathogenicity^67, 68^, and in yeast changes in the internal pH affect particle diffusion in the cytoplasm^29^. Here we investigate if surface attachment methods influence the internal pH of bacteria during time lapse imaging.

To monitor the effect of the adhesives on the internal pH of *E. coli* during growth, we use a genetically encoded indicator pHluorin^17, 69, 70^. pHluorin is a variant of the green fluorescent protein with pH sensitive spectrum that responds in a ratiometric manner, SI Figure 6. Prior to the growth experiments, pHluorin has been calibrated *in vivo* and *in vitro*, see *Supplementary Materials*^17^. For *in vivo* calibration we used various ∆*pH* = *pH*_*external*_ − *pH*_*internal*_ collapsing agents and noticed that the calibration curves deviate slightly depending on the uncoupler, which compromises the accuracy of the potential pH measurements. Though it is not clear what causes the difference in the calibration curves^17^, we show that the combination of potassium benzoate and methylamine hydrochloride (PBMH) allows us to reproduce the *in vitro* calibration most accurately (SI Figure 7), and we subsequently use PBMH for *in vivo* calibration.

Having calibrated pHluorin, we measure the intracellular pH of the immobilised bacteria during growth and division, and as before tracking the two generations (*G*_1_ and *G*_2_). Figure 3 shows that cytoplasmic pH of cells grown in MM9 decreases on all tested surfaces, dropping mainly in the first generation (SI Figure 8 shows cell to cell variation). In contrast, cytoplasmic pH of the cells growing in LB slightly increases over both generations as depicted in the SI Figure 9.

**Figure 3.**
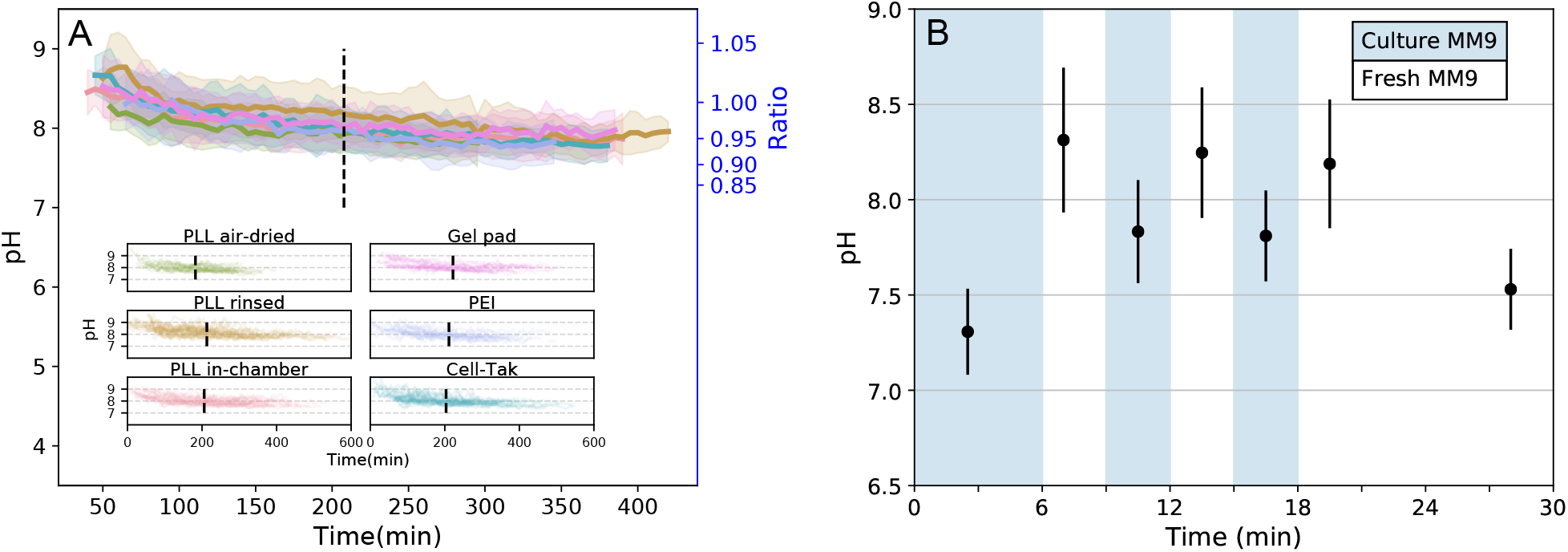
Intracellular pH dynamics of bacterial cells immobilised on a given surface. (**A**) Single cell intracellular pH is measured with cytoplasmic pHluorin during cell growth on different surfaces. Solid line and shaded area show the mean and standard deviation. Dashed line separates the mean first and second generations of bacteria. All tested immobilisation methods exhibit similar tendencies: pH values start from pH 8.38 ± 0.29 and decrease to about pH 7.78 ± 0.16 within ~7 h of observation. The inset shows all single-cell traces plotted for each condition. (**B**) Cells in a flow chamber are sequentially treated with the medium taken from a growing culture (blue regions) and fresh medium (white regions). Media exchange occurs in a short pulse manner at the beginning of each period, while no flow is applied during imaging. Data points show the mean of N *>* 500 measured bacteria with standard deviation as error bar.

The intracellular pH values we measured (Table 1) are higher than those commonly found in the literature (7.2-7.8^60, 61^). We assume these discrepancies originate from the fact that we constantly exchange the medium during the cell growth, removing from the environment metabolic waste products and any quorum sensing or signal molecules, which have previously been shown to influence cytoplasmic pH^62, 71, 72^. Indeed, when cells are kept in the original growth medium their cytoplasmic pH varies between 7.3 and 7.7, Figure 3B. It, however, increases rapidly to 8.2-8.3 when fresh medium is supplied and can be reduced back to *∼*7.8 upon the return to the original growth medium, Figure 3B. Further incubation in a fresh medium with no exchange (flow has been stopped) leads to the pH decrease to *∼* 7.5 after *∼* 10 min.

### Attachment quality on different surfaces varies

For single cell imaging it is important that analysed cells remain “flat”^18, 24^ (long axis parallel to the imaging plane) for the duration of observation, which could last several generations. We quantified the flatness by comparing the phase-contrast image intensity from two sides of the cells and defining a flatness score, Figure 4A (see *Materials and Methods* for details). All of the surfaces on which we obtained growth rates, “PLL in-chamber”, “PLL rinsed”, “PLL air-dried”, PEI and Cell-Tak show similar attachment quality during cell growth and division, Figure 4B. Interestingly, at the beginning of the observation even on APTES surface we find “flat” cells and correspondingly report a low flatness score in Figure 4B. Thus, grown in our media *E. coli* attaches to the APTES coated surface along the whole cell body, but it does so weakly (see SI Video 1). Images obtained on these surfaces with phase-contrast microscopy are indistinguishable, Figure 4C. In comparisons, cells on the gel pad surface are ‘flat’ as well, but we observe agarose impurities that influence image quality, Figure 4C.

**Figure 4.**
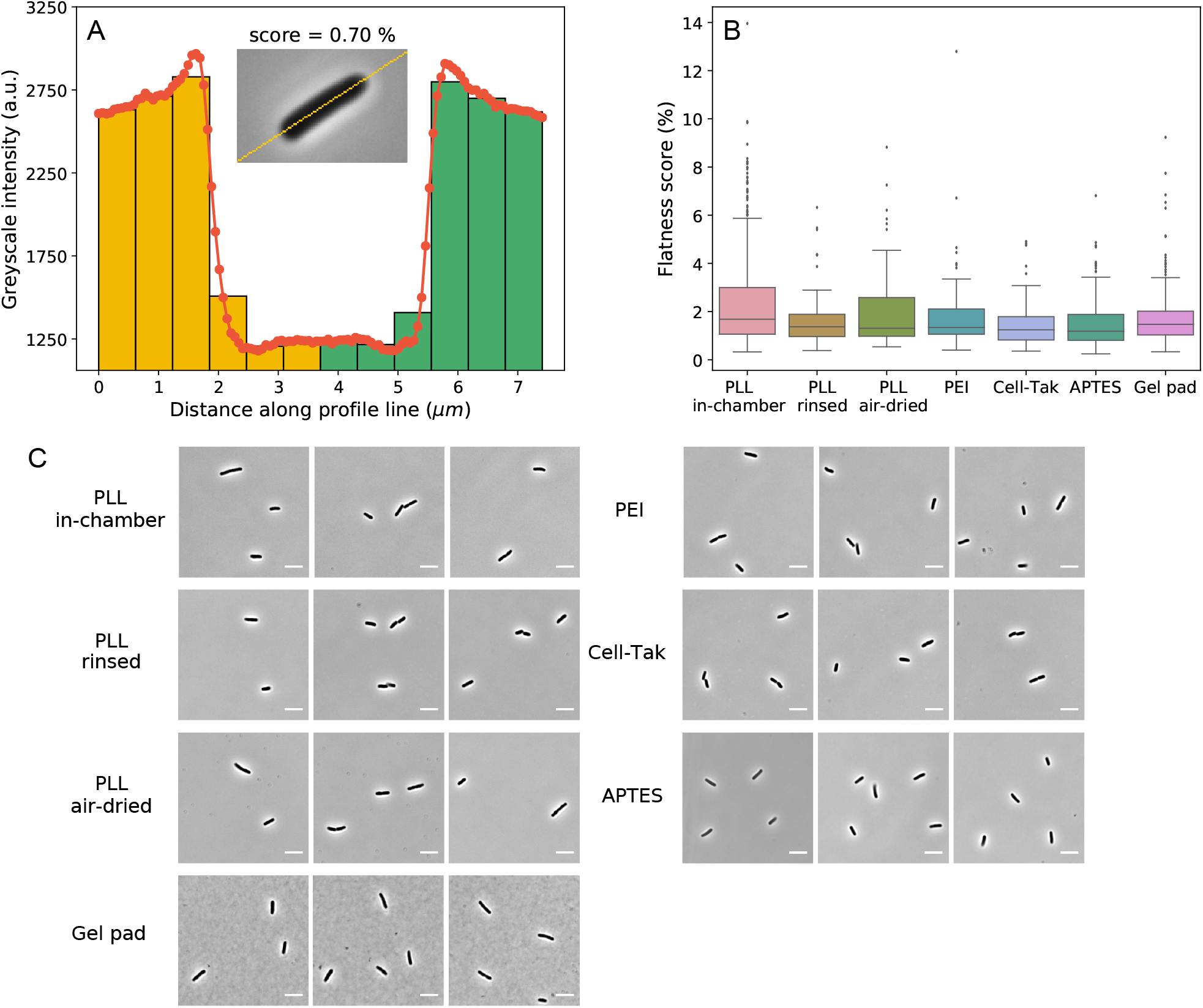
(**A**) The flatness score is calculated from the difference in the intensity between the two halves of the cell. Histogram of the intensity taken along the mid-line of the cell (inset) is shown. Two sides of the cells are indicated with yellow and green colour. Intensity of bins on the left and right that are equidistant from the middle are subtracted, and summed to obtained the flatness score. (**B**) Box plot (see *Materials and Methods* for details) of flatness scores on different surfaces obtained from 401, 51, 46, 61, 84, 143, and 269 cells for each of the surfaces (collected over ~ 130 µm^2^ flow chamber surface area), respectively. Here flatness score of 0% corresponds to a perfectly flat cell. (**C**) Examples of phase contrast images of *E. coli* cells attached to the surface with different immobilisation methods (scale bar is 5 µm). All images shows good “flatness” (see *Materials and Methods* for details) and contrast for cell morphology tracking.

## Discussion

Good surface attachment is an important requirement for bacterial single-cell studies using optical microscopy. However, we are unaware of a systematic study that characterises the effects on cells’ physiology caused by different adhesives. Changes in the cellular physiology caused by different surface attachment methods can influence not only studies of cellular physiology themselves, but also studies focusing on specific cellular molecular mechanisms. For example, metabolic rate or internal pH could lead to the alteration of cytoplasm properties, e.g. its fluidity^27, 29^, and many intracellular processes, including DNA replication and cell division, are highly dependent on the growth rate^73, 74^. It is, therefore, important to consider and characterise potential effects of the immobilisation method on physiology of the studied bacteria.

Here, we test a range of the immobilisation techniques and show that *E. coli*’s growth rate and shape are immobilisation method independent. Cell length and the growth rate are dependent on the growth medium, as expected, but independent of the surface attachment chemistry. We do, however, observe shortening of cells grow on the surface (Figure 2C and SI Figure 1 and 2), indicating an interesting physiological adaptation to growth on surface.

Though the concerns regarding use of PLL for surface attachment have been previously expressed in the literature^32, 75^, we note that the experiments that demonstrate inhibition of cell division by PLL, do so for the case of free PLL molecules in the medium^33^. We show that all three of the tested PLL-coating protocols leave no residual PLL in the medium and do not influence bacterial growth rate and accuracy of the division. It is possible that we see no adverse effects during growth on the PLL surface due to lower overal concentrations, or because the surface attached polymer does not integrate as efficiently into the *E. coli* membrane (insertion into the membrane has been reported for several different antimicrobial peptides^76^).

For measurements of *E. coli*’s cytoplasmic pH we use pHluorin and find that *in vivo* calibration curve is dependant on the agents used to collapse pH. We do not understand the observations at present, but speculate that it could occur due to the interaction between the uncoupler (CCCP or indole) and the bacterium, e.g. CCCP can be actively exported by EmrAB-TolC pump^77^. Using pHluorin we show that the internal pH of the attached *E. coli* is kept between 7.3 and 8.4 and doesn’t vary significantly with the surface coating. On all the tested surfaces in MM9 media the cytoplasmic pH decreases slightly in the course of the experiment, and the behaviour changes in LB medium, where cytoplasmic pH increases over the observation time. The result is not unexpected, as metabolic byproducts have been demonstrated to influence *E. coli*’s cytoplasmic pH^62, 71, 78–81^. For example, glucose metabolism mainly produces organic acids as byproducts, such as acetate, lactate, formate, succinate etc.^78^. These organic acids are capable of crossing the inner membrane in their uncharged form dissociating in the cytoplasm, which causes full or partial collapse of the pH gradient across the membrane^62, 71, 79^. In the case of LB medium, the alkalinisation of the media due to *E. coli*’s metabolism has been reported and attributed to the release of the amine-containing compounds^80, 81^. In both cases, however, the change does not exceed 0.6 pH units (an equivalent of 35.5 mV of proton motive force).

We were able to quantify growth in all but one tested surface. On APTES surface we observed attachment and growth, but the attachment was not strong enough to support growth for sufficient amount of time needed to quantify it. Here we note that our flow rate of 4 *µ*l/min was chosen to ensure a continuous supply of fresh oxygen and nutrients throughout our measurements. The average size of our flow chamber used for surface attachment is 17.5 *µ*l, which means the media is exchanged in ~ 5 min. If we take into account *E. coli’s* oxygen consumption rates^82, 83^ and the concentration of cells during imaging (calculated based on observed surface density), we obtain that in our flow chamber oxygen will be depleted in 14 min to 3.7 h. Thus, flow rate could have been reduced (in particular for conditions in which oxygen consumption rate is lower) potentially allowing growth on APTES surfaces as well.

In summary, we conclude that all the tested immobilisation protocols can be used for live cell imaging of *E. coli* without specifically affecting cells’ main physiological traits.

## Materials and Methods

### Bacterial Strains and Growth Conditions

The strain, EK03, is the *Escherichia coli* K-12 MG1655 strain with “sticky” flagella mutation^17, 84^ and pkk223-3/pHluorin(M153R) plasmid. The plasmid containing pHluorin with M153R mutation to have better stability of fusion proteins^69^ was a kind gift from Dr. Tohru Minamino. SI Figure 5 shows growth comparison of the two strains, EK03 and wild-type MG1655, in Lysogeny broth (LB: 1% w/v Bacto-Tryptone, 0.5% w/v yeast extract and 0.5% w/v NaCl, pH 6.8) and MM9 medium (Na_2_HPO_4_ 50 mM, NaH_2_PO_4_ 20 mM, NaCl 8.5 mM, NH_4_Cl 20 mM, CaCl_2_ 0.1 mM, KCl 1 mM, MgSO_4_ 2 mM, 0.5% glucose, and MEM Amino acides solution (Gibco, USA), pH 7.0) at 37°C and 23°C shaken at 220 rpm. Cells were diluted from an overnight culture to achieve steady state growth (6 to 10 divisions before reaching OD ~ 0.4).

For imaging cells were inoculated in LB or MM9 medium from an overnight culture grown in LB. Cultures are grown for 2 h in LB and 4 h in MM9 meidum at 37°C with continuous shaking (220 rpm) to reach optical density of ~ 0.4 (achieving ~ 3 divisions). Before imaging cells were further diluted 1:60 into MM9 or LB for the surface immobilisation protocols. For comparison of our growth rates at different temperatures and in different media please see SI Table 2.

### Microscope and Microfluidic chambers

Bacterial growth assays were conducted using a motorised, inverted optical microscope (Ti-E, Nikon, Japan) with perfect focus system for time lapse observation. The microscope is equipped with a 100x Objective (Plan Apo 100x/1.45NA lambda, Nikon, Japan), sCMOS Camera (Zyla 4.2, Andor, UK) and LED fluorescent excitation light source (PE4000, CoolLED, UK). Imaging was performed in phase-contrast and epifluorescence configuration, the latter was used for measuring the cytoplasmic pH with pHluorin. The exposure times for phase-contrast and epifluorescence imaging were 100 and 70 ms respectively, and images were recorded every 5 min. The pH measurement using pHluorin was achieved by ratiometric method from two excitation wavelength of 395 and 470 nm and emission of 510 nm images. The excitation and emission was achieved by LED excitation light (395 nm and 479 nm) via a dual-band dichroic mirror (403/502nm, FF403/502-Di01-25×36, Semrock, USA) and a dual-band bandpass emission filter (FF01-433/530-25, Semrock, USA).

To supply fresh oxygenated media throughout the experiment we use a flow chamber made as follows. Two 1.5 mm holes were drilled in a microscope slide 20 mm apart. PTFE tubing with inner diameter 0.96 mm was attached to the slide with epoxy glue, SI Figure 10. The flow chamber was then created by attaching double sided tape or gene frame (Fisher Scientific, USA) to the slide and covering it with pre-coated or uncoated cover glass depending on the immobilisation protocol. Gene frame was used to create a larger chamber to fit the agarose pad, while sticky tape was used for all of the other protocols. Dimensions of the formed flow chamber are 3.5 × 25 × 0.2 mm for doubled sided tape, and 17 × 28 × 0.25 mm for gene frame. The flow chamber construction protocol varied slightly with different coating protocols. For the PLL “in chamber” and Cell-Tak coating methods, the flow chambers were sealed before coating. In other cases, the coverslips were coated prior to the flow chamber construction.

For all of the immobilisation assays medium was flown at 400 µl/min flow rate at the end of the attachment protocol for 4 min to remove poorly attached cells, upon which the flow rate was altered to 4 µl/min for the duration of the experiment (12 h). Media was flown with a syringe pump (Fusion 4000, Chemyx, USA). Before the imaging begun we identified a field of view with maximum number of “flat” cells^18, 24^ for each of the surfaces, and the flatness score (see below for further details) is calculated on thus selected fields of view.

### Immobilisation protocols

#### Preparation step: coverslip cleaning

The coverslip is sonicated in an ultrasonic bath (D150H, Delta, Taiwan) with saturated solution of KOH in ethanol for 30 min. It is then rinsed with the deionised water and sonicated for further 30 min in deionised water. Cell do not attach to so treated glass without suitable coating. The cleaning step has been performed prior to all attachment protocols.

#### PLL “in-chamber”

Surface of the flow chamber is coated with 0.1% poly-L-lysine (PLL) by flushing PLL through the channel for ~ 10 s followed by washing it out with the excessive volume of growth medium (x20 times the volume of the chamber). Cells are then loaded into the flow chamber and incubated for 1 to 3 min to allow attachment, and then washed out as described under *Microscopy and Microfluidic chambers* section. The total length of time it took to prepare a flow chamber with cells attached is ~ 40 min, irrespective of the attachment method.

#### PLL “rinsed”

Coverslip is coated with the PLL prior to the flow chamber construction. 100 µl of the 0.01% PLL solution (diluted from P8920 Sigma-Aldrich, USA) is spread over approximate 1.5 cm^2^ area. The solution is allowed to sit on the coverslip for 30 min, then washed off with 5 ml of deionised water. The coated coverslip is then used to construct a flow chamber. The cells are attached as above.

#### PLL “air-dried”

The protocol is similar to the “rinsed” method. Here, the PLL solution on the coverslip is air dried fully, typically for 1.5 h at room temperature. The coated coverslip is then washed with 5 ml deionised water and used to construct a flow chamber. The cells are attached as above.

#### PEI

200 µl of 1% PEI is spread out on the coverslip covering the area the size of the flow chamber tunnel. Solution is incubated on the surface for 10 s, and washed off thoroughly with 100 ml of deionised water. The volume of the water should be much higher than that of the PEI to leave no residual PEI molecules that are not attached to the glass surface. Otherwise (e.g. if we use 5 ml of water), we observe cells blebbing^85, 86^ and dying when grown on the surface. We also notice that the “in-chamber” coating method is not applicable for PEI in MM9 media, as it leads to MM9 precipitation in the chamber caused by the leftover free PEI molecules. This is not the case for LB; “in-chamber” coating method can be used with PEI and LB^18^. The cells are subsequently attached as described above.

#### Cell-Tak

Cell-Tak working mixture is prepared by adding 14 µl of Cell-Tak (1.16 mg/mL) to 174 µl NaHCO_2_ (pH 8.0) followed by immediate vortexing. A pre-assembled flow chamber is incubated with Cell-Tak mixture for 20 min, then washed with 3 ml of deionised water. 3 ml of MM9 is flushed through the chamber before attaching the cells. Finally, cells are attached as above.

#### APTES

A coverslip was incubated in 2% APTES for 2 h, and then rinsed with 5 ml deionised water and 5 ml acetone. The remaining acetone was air-dried with nitrogen. The coated coverslip was later used for the flow chamber construction. Cells are attached as above.

#### Agarose Gel Pad

For the agarose gel pad, the flow chamber area was 17 × 28 mm^2^. The gel pad was created by adding a 5 µl droplet of melted 1% agarose to the middle of the flow chamber. 0.5 µl of the cell culture was added onto the solidified pad and covered immediately with the coverslip. The fresh medium was constantly circulated around the agarose “island” during the experiment.

### Image analysis

#### Cell segmentation

Phase contrast images of the cells were analysed with custom written Python script and ImageJ^87^. In phase-contrast microscopy, cells appear as dark objects on a light background, with a characteristic white halo, Figure 1 inset. Cells are segmented with Watershed algorithm^88, 89^ implemented as ImageJ Marker-controlled Watershed plugin^90^. The algorithm treats an image as a topological surface where the pixel intensity corresponds to the area height. It then identifies the edge of the cell as the watershed separating neighboring drainage basins. The limit of watershed algorithm grows based on the background intensity. After the segmentation, the cell length is calculated by PSICIC algorithm (Projected System of Internal Coordinates from Interpolated Contours)^91^. Briefly, the algorithm finds two poles of a cell as points that are the greatest Euclidean distance apart, thus creating two curves. On each of the two contour curves the algorithm evenly distributes equal number of points and then connects them (effectively creating width lines). Finally, the center line, i.e. the length of the cell, runs along the middle of the width lines^91^. We define the cell width as the average of all the width lines which are greater than nine-tenths of the maximum width line. The division time is determined manually by analyzing the cells from the onset of constriction. It is difficult to set the criteria for an exact division time point based on the morphology or the gray scale intensity, because the cells can tilt (in the x and y plane) during the division. The manual criteria are, for example, the detachment of one of the daughter cells (we often find that the daughter cell detaches due to flow), or centre lines of the two cells might no longer overlap. Subjectivity in setting a division time point can cause time error of about ± 10 min, which affects division time measurements but not the estimates of the growth rate *b*.

#### Flatness score

To quantify the quality of attachment on a given surface we define a flatness score. We start by extending the center line running along the middle of the cell to include cell background and split it into 12 points to obtain a histogram of intensity along the extended centre line. The height of each histogram bin is the average grey scale value of pixels whose positions are located within the interval of the bin. Next, the 12-bin histogram is normalised to 1. Lastly, values of the bins on each side of the cell that are equidistant from the middle are subtracted and summed to obtain the flatness score. A perfectly flat cell should have a symmetric profile and a flatness score of zero.

### Numerical and statistical method

#### Growth curve fitting

The growth of the cell follows *L*(*t*) = *L*_*b*_ × *e*^*bt*^, where *L*_*b*_ is the cell initial length, *t* time in minutes, *b* the growth rate^46^. The equation is fitted to the cell length using Levenberg-Marquardt algorithm^92^, a non-linear least squares method, implemented using Scipy, a numerical package in Python^93^. Initial parameters we identified by first fitting a polynomial to the logarithmic growth of the cell length.

#### Box plot

The borders of the box and the middle line in our box plots indicate the three quartiles of the data. The range of whiskers is 1.5 of the interquartile range, and data outside the whiskers are the outliers (displayed as dots).

## Acknowledgements

This project is supported by the Human Frontier Science Program Grant (RGP0041/2015) to TP, CJL and FB. CJL is financially supported by the Ministry of Science and Technology, Republic of China under contract No. MOST-107-2112-M-008-025-MY3. EK was supported by the Global Research and Principal’s Career Development PhD Scholarships. We thank Tom Shimizu and Victor Caldas (AMOLF) for sharing the APTES protocol of immobilization.

## Author contributions statement

YKW, EK, FB, TP and CJL designed research. YKW, SYL and EK performed research and analysed data. YKW, SYL, EK, FB, TP and CJL interpreted results and wrote the paper. All authors reviewed the manuscript.

## Supplementary Materials

### Supplementary Methods: pHluorin calibration

The *in vivo* calibration of pH sensor was performed as follows^17^. The mixture of 100 mM MES, HEPES and AMPSO buffers was adjusted to a set of pH values in the range between 5.5 and 9, and supplemented with one of the three pH collapsing agents: 40 mM potassium benzoate and 40 mM methylamine hydrochloride (PBMH)^65^, 25 µM CCCP or 5 mM indole^94^. Tunnel-slides were prepared as previously described^17, 70^: two bits of sticky tape form a tunnel and are sandwiched between a coverslip and a microscope slide. Buffer of known pH was flushed into a channel, incubated for 15 min, upon which 5 different fields of view containing over 100 cells were imaged with 50 ms exposure time. The calibration curves were plotted as ratio of emission intensities for excitation at 395 nm and 475 nm against pH, and fitted with the sigmoid function 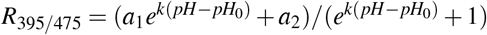, where *a*_1_, *a*_2_, *k* and *pH*_0_ are free fitting parameters.

*In vitro* calibration was performed with the purified pHluorin protein diluted into buffer of known pH in the 96-well plate (Thermo Scientific, Optical bottom). The pHluorin excitation spectra for 510 nm emission was measured in Spark 10M multimode plate reader (Tecan Trading AG, Switzerland). The His-tagged protein was purified using affinity chromatography column^17^. The excitation spectra was scanned from 380 nm to 480 nm with 5 nm step size. Additionally, the autofluorescence of the buffer with no added protein was measured and subtracted from the measured protein intensity.

### Supplementary Videos

SI-Video 1: Time lapse movie of cells growth on the APTES surface, total time 10 hours.

SI-Video 2: Time lapse movie of cells growth on the PLL in-chamber surface, total time 12 hours.

SI-Video 3: Time lapse movie of cells growth on the PLL rinsed surface, total time 12 hours.

SI-Video 4: Time lapse movie of cells growth on the PLL air-dried surface, total time 12 hours.

SI-Video 5: Time lapse movie of cells growth on the PEI surface, total time 12 hours.

SI-Video 6: Time lapse movie of cells growth on the Cell-Tak surface, total time 12 hours.

SI-Video 7: Time lapse movie of cells growth on the Gel pad surface, total time 12 hours.

---

**Table 2.** Population doubling times at different temperatures (*t*_*D*_ = *ln*(2)*/λ*). Nutrient Broth (Meat extract +0.2% peptone +0.16% glucose and Salt solution), 199 Tissue Culture Medium (as in^95^), 0.2% Glucose Medium (0.2% glucose + Salt Solution), 0.2% Succinate Medium (0.2% succinate + Salt Solution), 0.2% Lactate Medium (0.2% lactate + Salt Solution), Salt Solution (MgSO_4_ ⋅ 7H_2_O – 0.1, citric acid – 1.0, Na_2_HPO_4_ ⋅ 2H_2_O – 5.0, Na(NH_4_)HPO_4_ ⋅ 4H_2_O – 1.74, KCl – 0.74 g/l), Minimal Medimum (defined MOPS medium^96^), Rich Glucose Medium (MOPS medium supplemented with 0.4% glucose, amino acids (minus leucine; 0.12 mM valine and 0.08 mM isoleucine), 0.01 mM of each of the five vitamins (p-aminobenzoic acid, p-dihydroxybenzoic acid, p-hydroxybenzoic acid, pantothenate (calcium salt), and thiamine) and 0.2 mM of each of four bases (adenine, guanine, cytosine, and uracil)).

| Media and Strain | $t_D$ at 37°C | $t_D$ at 25°C | $t_D$ at 23°C | Reference |
| --- | --- | --- | --- | --- |
| Nutrient Broth, *Salmonella typhimurium* | 25 min | 56 min | — | 47 |
| 199 Tissue Culture Medium, *Salmonella typhimurium* | 32 min | 68 min | — | 47 |
| 0.2% Glucose Medium, *Salmonella typhimurium* | 50 min | 92 min | — | 47 |
| 0.2% Succinate Medium, *Salmonella typhimurium* | 65 min | 125 min | — | 47 |
| 0.2% Lactate Medium, *Salmonella typhimurium* | 67 min | 120 min | — | 47 |
| Rich Glucose Media, *E. coli* B/r NC3 | 33 min | — | 100 min | 48 |
| Minimal Medium, *E. coli* B/r NC3 | — | — | 157 min | 48 |
| LB, *E. coli* K-12 MG1655 | 22 min | — | — | This work |
| MM9 Medium, *E. coli* K-12 MG1655 | 49 min | — | — | This work |
| LB, *E. coli* K-12 EK03 | 26 min | — | 86 min | This work |
| MM9 Medium, *E. coli* K-12 EK03 | 58 min | — | 210 min | This work |

**SI Figure 1.**
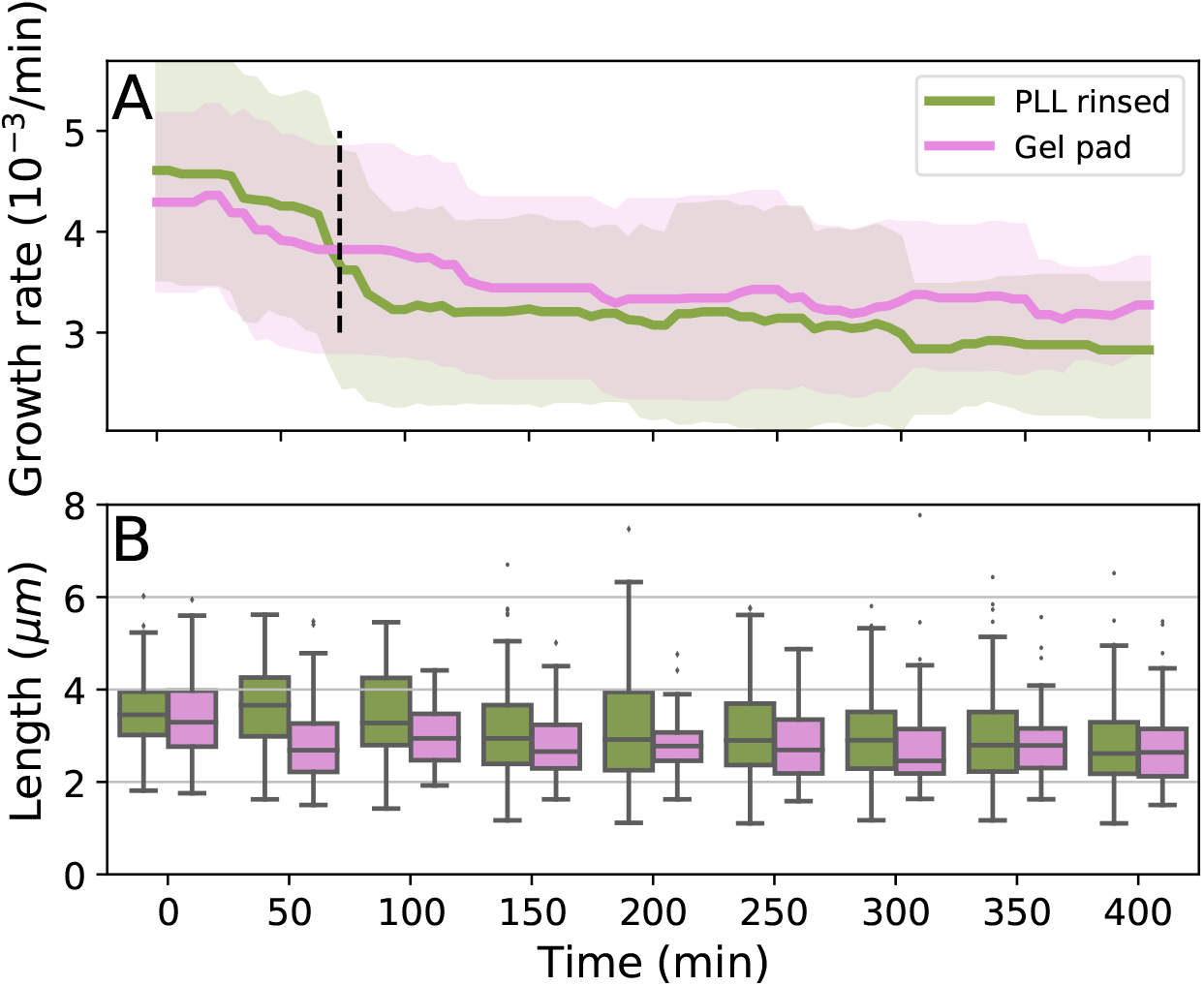
(**A**) Growth rate (*b*) decreases on PLL rinsed and gel pad surfaces during *G*_0_ (see Figure 1, top). Dotted horizontal line at 73 min indicates completion of *G*_0_ and beginning of *G*_1_, after which the growth rate remains roughly constant. We attribute the reduction in growth rate during *G*_0_ to adaptation to growth at 23°C after the culture has been grown at 37°C (see also *Materials and Methods*. (**B**) Cell length distribution over time on the PLL rinsed and a gel pad surfaces changes in line with growth rate changes in (A). The average length of the cells population decreases from 3.56 ± 0.81 µm to 2.75 ± 0.80 µm on the PLL rinsed, and from 3.41 ± 0.78 µm to 2.73 ± 0.81 µm on the gel pad. A steady length distribution is reached by *G*_1_. Cells grow on the other tested surface showed similar length distribution over time.

**SI Figure 2.**
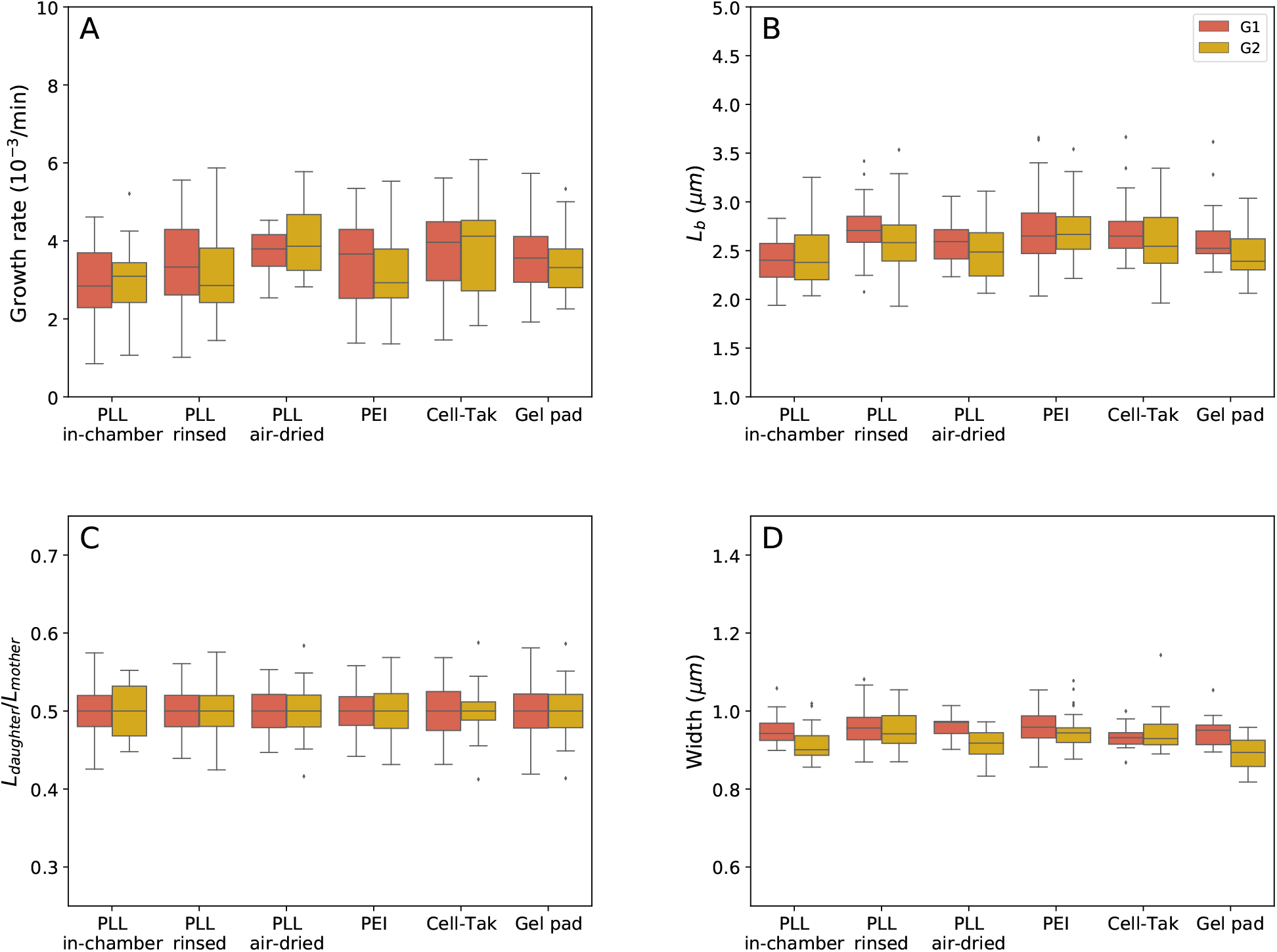
Comparison of (**A**) the growth rate, (**B**) initial cell length *L*_*b*_, **(C)** length ratio *L*_*daughter*_/*L*_*mother*_ and **(D)** cell width for G1 and G2 (see Figure 1, top).

**SI Figure 3.**
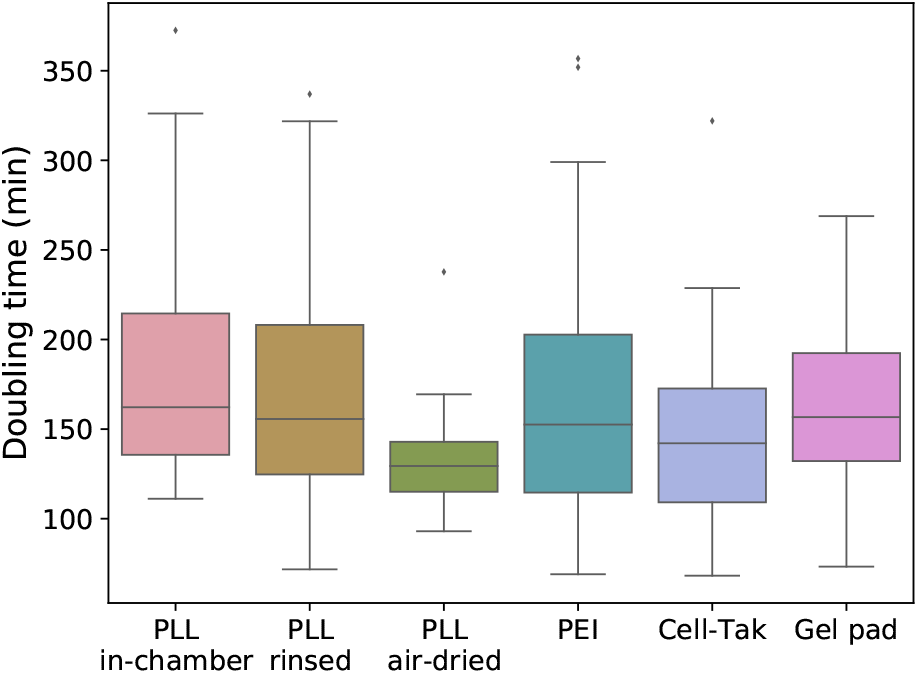
Doubling time *t*_*d*_ = ln (*L*_*d*_/*L*_*b*_)*/b*, where *b* is the individual cell’s growth rate.

**SI Figure 4.**
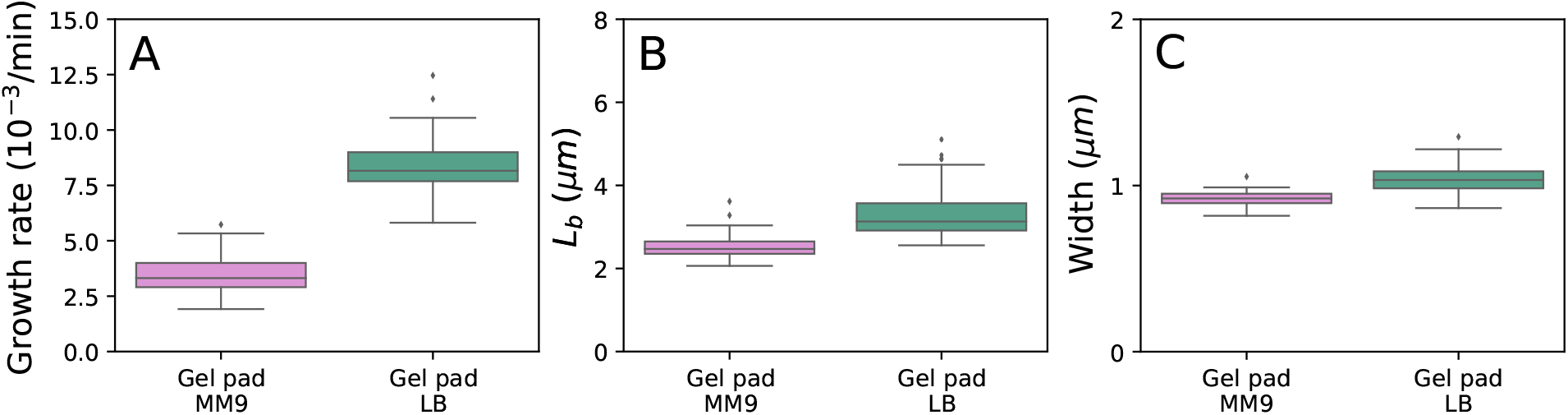
Comparison of (**A**) the growth rate (*b*), (**B**) initial cell length *L*_*b*_ and **(C)** cell width for cells growing on the gel pad in MM9 and LB media.

**SI Figure 5.**
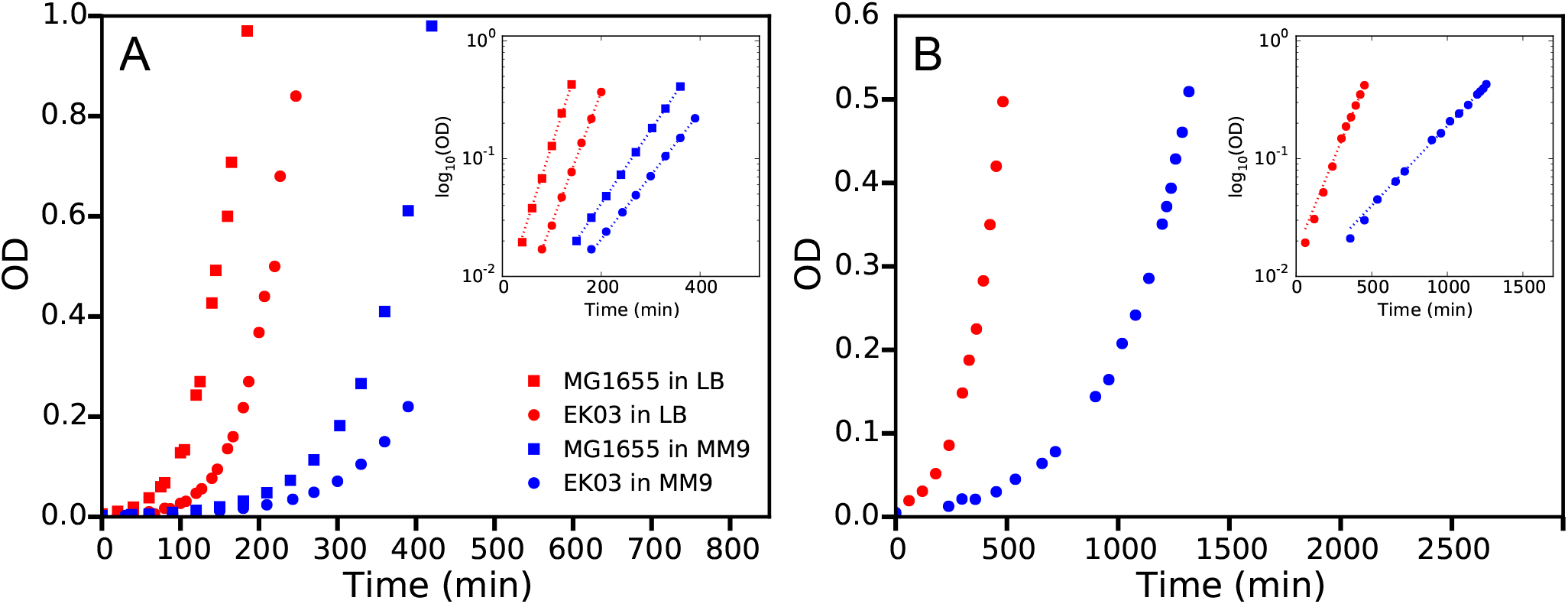
Growth curves of MG1655 (squares) and EK03 (circles) strains grown in LB (red) or MM9 (blue) media in the flask at 37 °C (**A**) and 23 °C (**B**) from 10^*−*5^ starting dilution of the overnight culture. Time scale is renormalised to exclude lag times from the graph, i.e. renormalised *t* = 0 is the time of lowest measurable OD value. Inset: log_10_(OD) is plotted against time to calculate the growth rates (*λ*). Dotted lines show exponential fits for OD values between 0.02 and 0.4. See calculated growth rates in SI Table 2.

**SI Figure 6.**
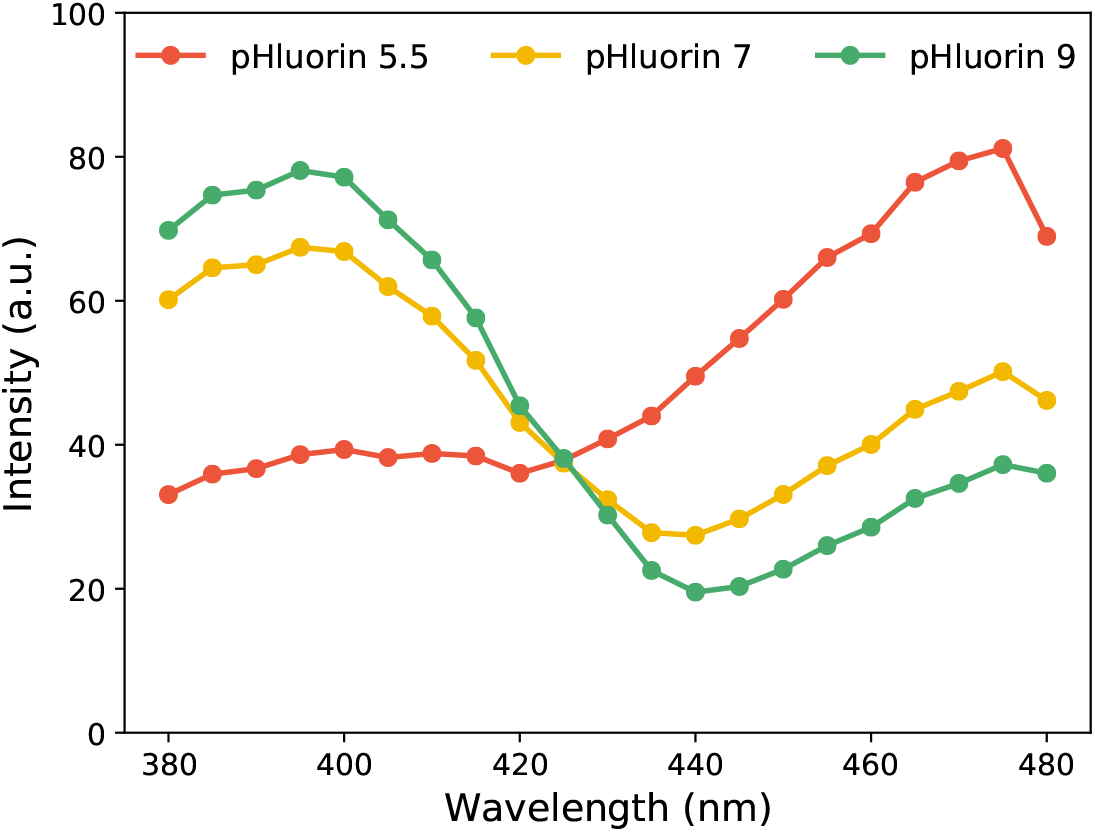
Excitation spectra of purified pHluorin at pH values 5.5 (red), 7.0 (yellow) and 9.0 (green). Emission is collected at 510 nm.

**SI Figure 7.**
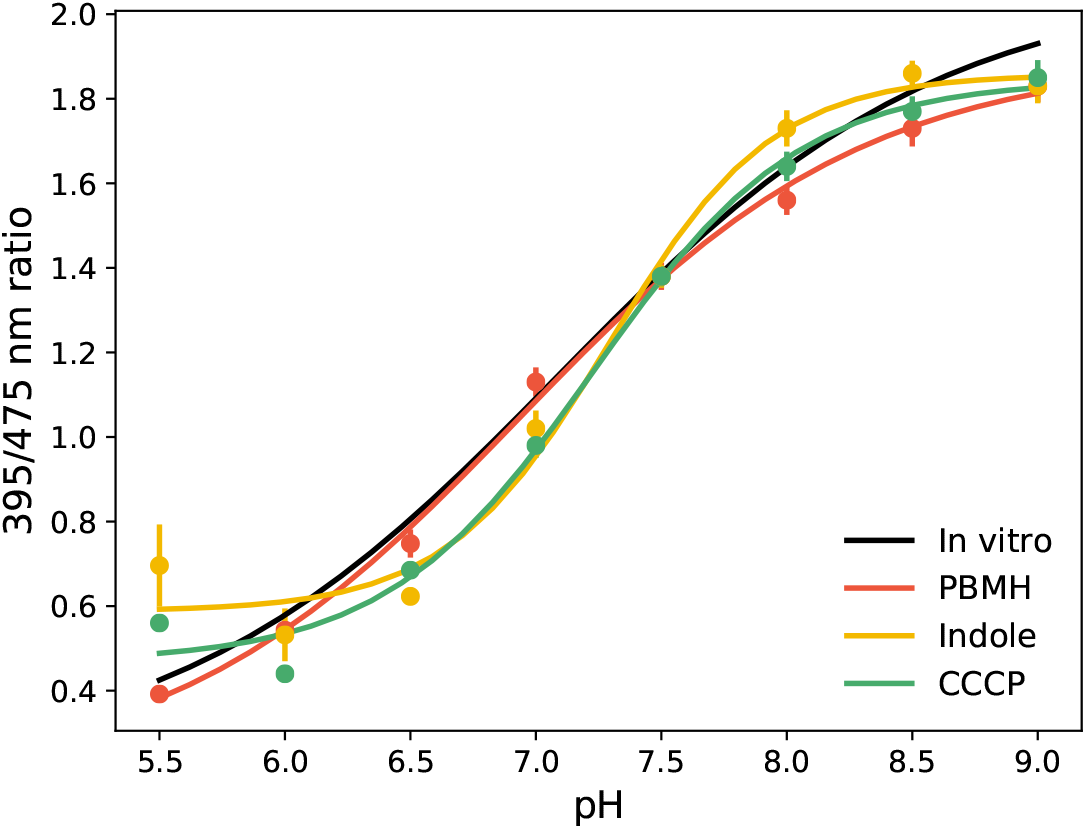
Comparison of the *in vivo* and *in vitro* calibration curves. Coloured lines show *in vivo* pHluorin calibration curves with 40 mM PBMH (red), 5 mM indole (yellow) or 25 *µ*M CCCP (green) as ∆pH collapsing agents. Black curve shows *in vitro* calibration curve of purified pHluorin in buffer with no supplements. *In vivo* curve with PMBH aligns best with the *in vitro* curve.

**SI Figure 8.**
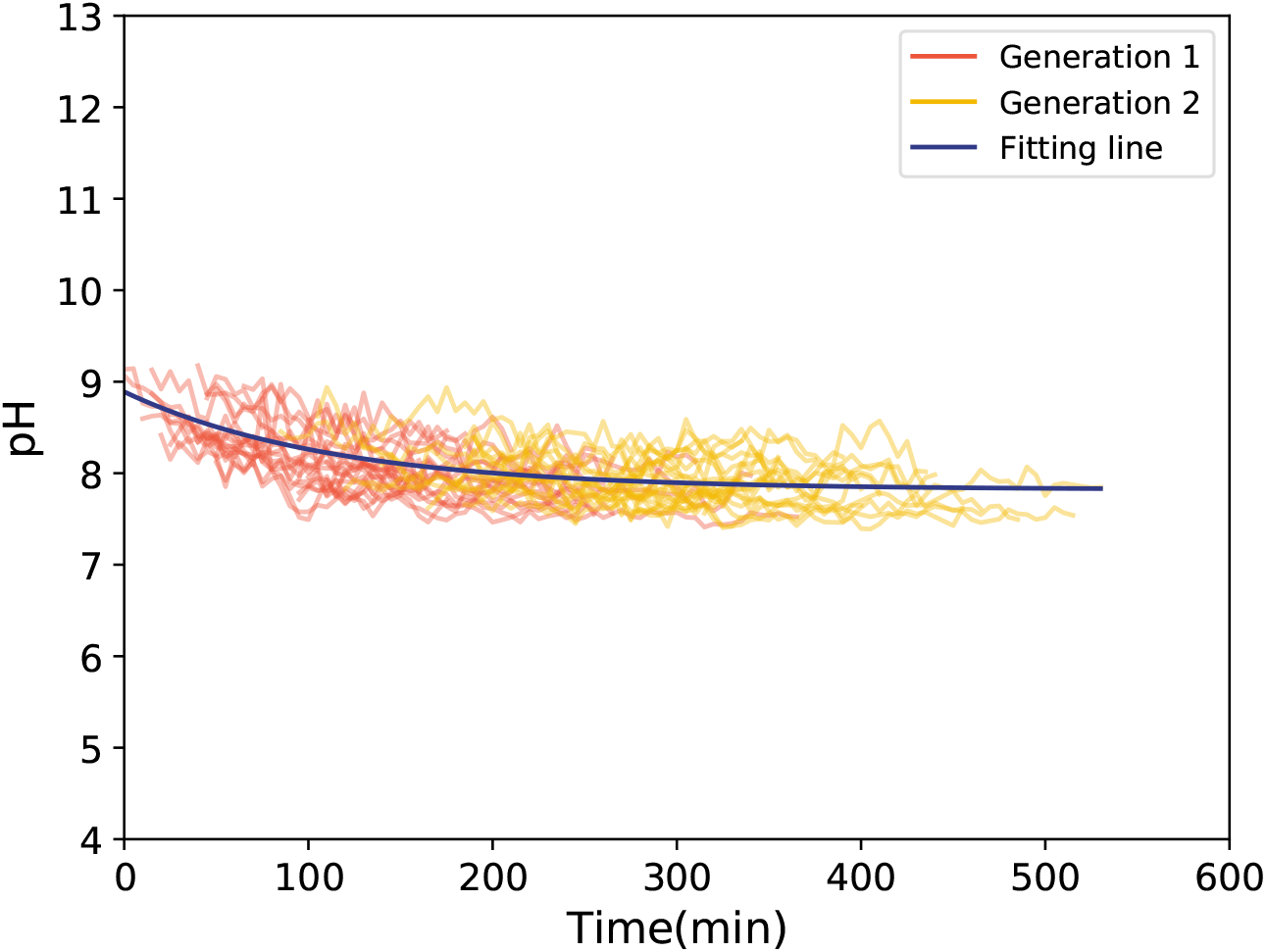
Single cell intracellular pH dynamics for two generations of bacteria, first generation is shown in red and the second in yellow. Cells are grown on the PLL “in-chamber” coated surface. An exponential, shown in blue, was fitted to the whole data set to obtain the final value of pH. The single cell intracellular pH on other coated surfaces shows similar decay.

**SI Figure 9.**
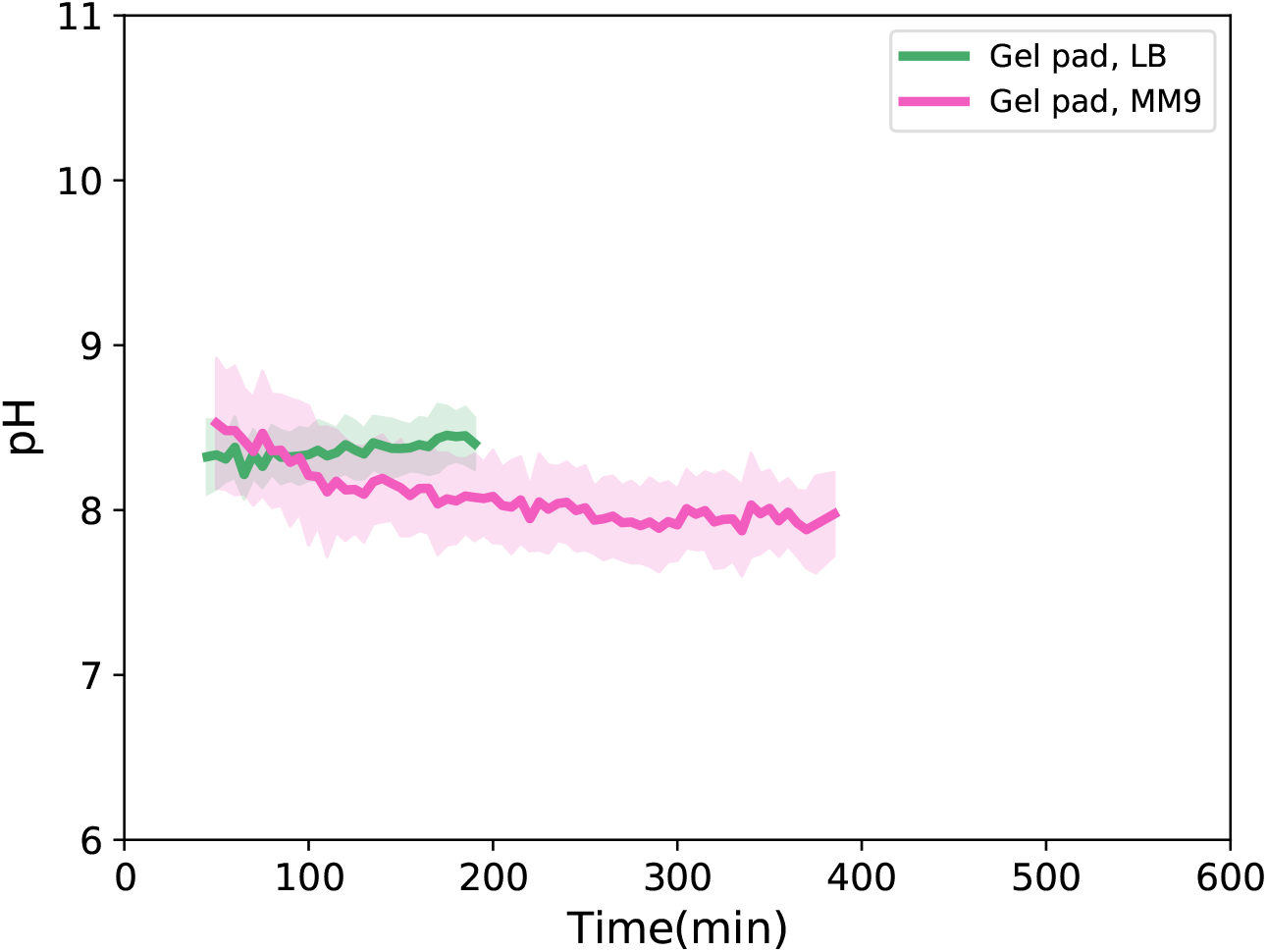
Mean and the standard deviation of the intracellular pH of the cells grown in the gel pad in LB (green) or MM9 (red) media.

**SI Figure 10.**
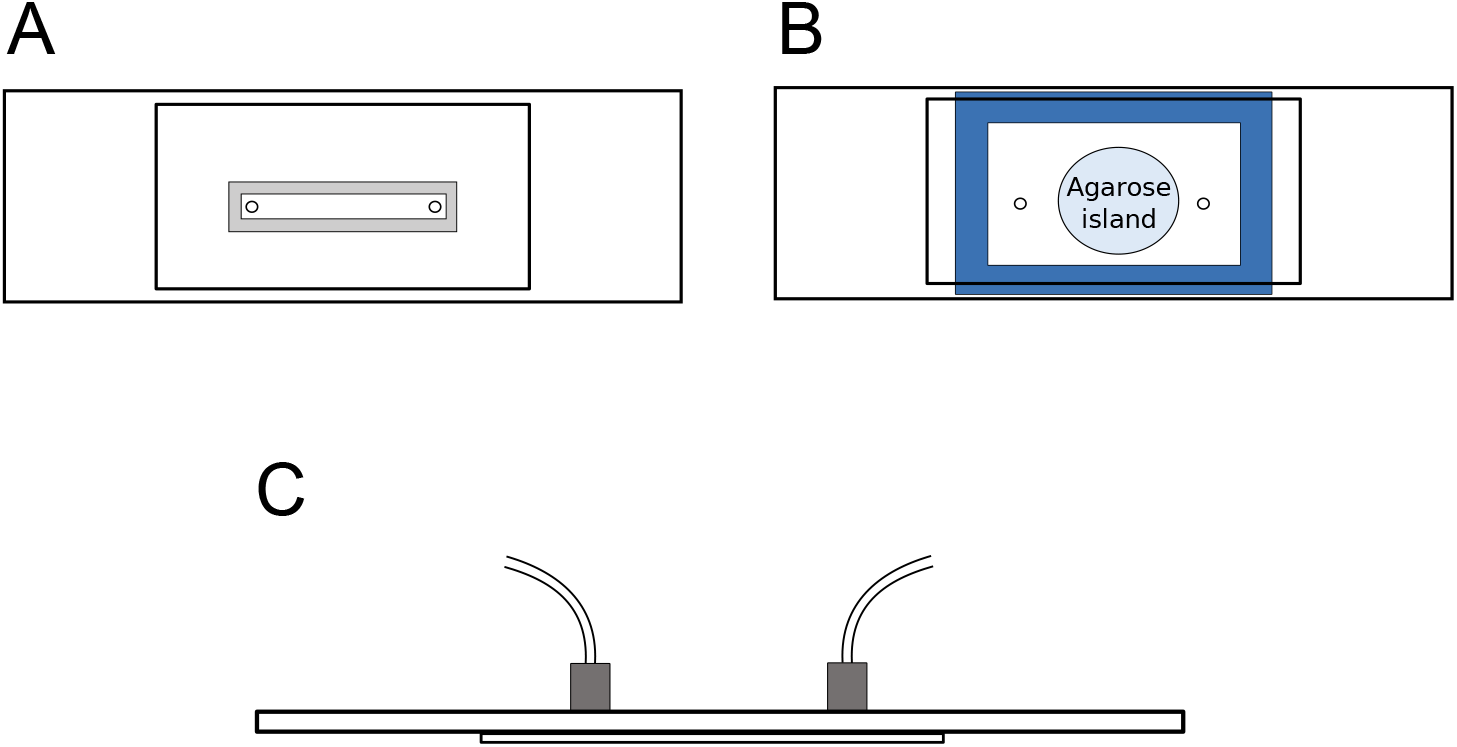
Schematic of our flow chambers for bacterial immobilsation experiments. (**A**) top view of the flow chamber used for surface attachments, (**B**) top view of chamber for agarose island experiments, and (**C**) side view of flow chambers.

## Notes

#### Summary of Updates

We have included additional analysis of the data as well as an additional cartoon explaining the time sequence of the experiment.

